# Secondary DNA transfer on denim using a human blood analogue

**DOI:** 10.1101/2021.11.25.470033

**Authors:** Rebecca Ridings, Alon Gabriel, Colin I. Elliott, Aaron B.A. Shafer

## Abstract

DNA quantification technology has increased in accuracy and sensitivity, now allowing for detection and profiling of trace DNA. Secondary DNA transfer occurs when DNA is deposited via an intermediary source (e.g. clothing, tools, utensils). Multiple courtrooms have now seen secondary transfer introduced as an explanation for DNA being present at a crime scene, but sparse experimental studies mean expert opinions are often limited. Here, we used bovine blood and indigo denim substrates to quantify the amount of secondary DNA transfer and quality of STRs under three different physical contact scenarios: passive, pressure, and friction. We showed that the DNA transfer was highest under a friction scenario, followed by pressure and passive treatments. The STR profiles showed a similar, albeit less pronounced trend, with correctly scored alleles and genotype completeness being highest under a friction scenario, followed by pressure and passive. DNA on the primary substrate showed a decrease in concentration and genotype completeness both immediately and at 24 hours, suggestive of a loss of DNA during the primary transfer. The majority of secondary transfer samples amplified less than 50% of STR loci regardless of contact type. This study showed that while DNA transfer is common between denim, this is not manifested in full STR profiles. We discuss the possible technical solutions to partial profiles from trace DNA, and more broadly the ubiquity of secondary DNA transfer.

## 1. Introduction

Technology has enabled the detecting and profiling of minute amounts of DNA [1], yet it is still difficult to reliably attribute small amounts of DNA to a biological source [2]. The lack of biological context complicates the understanding of the events that led to the deposition of DNA (e.g. [3]), and is broadly referred to as trace DNA. Advancements in diagnostic markers are starting to become useful for attributing trace RNA and DNA to biological sources [4], but the temporal window is likely limited [5]. Secondary DNA transfer occurs when DNA-containing materials (e.g. skin-cells, blood, saliva, semen) are deposited on a surface, and the material (or DNA) is subsequently transferred to another surface upon contact [6]. Defense counsel have now argued that secondary DNA transfer could explain why DNA was present at crime scenes or on a piece of evidence [6].

Secondary transfer of DNA can occur through a variety of means: for example, contamination at both the crime scene and lab can lead to transfer [7–9], with gloves being particularly problematic [10]. Other more benign mechanisms of DNA transfer include shared laundry [11] and basic social interactions [12]. Numerous reviews have placed a spotlight on the issue and often cite real cases [6,8,13–15], but the individuality of each scenario hampers generalities when it comes to the nature of secondary DNA transfer. Meakin and Jamieson [6] suggested it is impossible to reliably infer the mode of transfer, while Champod [14] had a more nuanced take, suggesting a balance between data and personal experience could inform opinions. Importantly, Champod [14] argued that experience was not a substitute for the systematic acquisition of data.

Controlled experiments offer baseline data that can help inform an expert opinion. For example, when two surfaces touch we now know that DNA transfer is dependent, in part, on substrate material [12], length of contact time [16], and individual (shedder) status [17]. When for example, a cotton object contains evidentiary DNA, it is relevant to know that blood (and hence DNA) are more readily transferred to cotton than wool and plastic substrates [12]. We were particularly interested in the transfer of DNA between denim, a cotton textile, given the global ubiquity and popularity of the material [18]. Denim has long history as evidence, notably as it pertains to fiber analysis [19], but extracting and analyzing DNA from biological material on denim is also common. The use of indigo dye can be problematic for detection of dye-based assays like qPCR [20] or hamper DNA polymerase activity [21]. Commercial DNA extraction and PCR kits remove such inhibitors and show high genotyping success as a result [22], while qPCR interference requires large and unrealistic volumes on indigo dye [20]. Here, we adapt the experiment of Goray et al. [12] to quantify both the amount of DNA and quality of STRs from a transfer event under three contact scenarios between indigo denim.

## 2. Materials and methods

### 2.1 Sample preparation

Whole bovine blood was collected from individual cows at a local abattoir. We initially collected blood without anticoagulant; but to avoid immediate clotting we also conducted experiments where acid citrate dextrose anticoagulant solution A (ACD-A) was added to the bovine blood in 12.5% v/v upon collection. The ACD-A solution was prepared by dissolving 6.6 grams of sodium citric dihydrate, 2.4 grams of citric acid and 6.68 grams of dextrose anhydrous, purchased from ACP Chemicals Inc. The blood was placed on ice for transportation back to the laboratory (^~^15 minutes). Whole mammalian blood with the inclusion of ACD-A is an appropriate substitute for studies of this nature [23–25] as bovine blood exhibits similar fluid properties to that of humans [26]. We compared the DNA transfer concentrations, with and without ACD-A using a Two-sample T-test, to test for an effect.

### 2.2. Simulated transfer and DNA extraction

Indigo denim fabric acted as both the primary and secondary substrates in the simulated transfer events. The denim was cut into 2×2 cm squares and autoclaved using the pre-vacuum 1 setting (121.1°C for 30 mins, with a drying time of 15 mins) of the Getinge 533LS Sterilizer. A total of 25 μL of fresh bovine blood was deposited onto the middle of the primary substrate. This volume of blood was chosen because blood contains ^~^20 ng of DNA per μL, meaning if only 1% of the total DNA were transferred it would still result in ^~^5 ng of DNA [12]. Ten minutes after the initial deposition of blood, one of three secondary transfer events was simulated using an approach modeled after Goray et al. [12]: passive, pressure, or friction contact (Figure 1). Passive transfer was conducted by placing the primary substrate (A) onto the secondary substrate (B) for 60 seconds. The pressure transfer event was simulated using a ^~^160 g weight placed on top of the two substrates, again with A laying on top of B for 60 seconds. The transfer via friction had substrate A laid on top of B and subsequently placed on a plate vortexer, which shook at a speed of 2000 rpm for 60 seconds. A minimum of ten technical replicates of substrate B were completed for each contact type and the samples were left to dry overnight at room temperature (^~^22°C and 50% relative humidity). Denim with no blood acted as a negative control for the extraction. Whole blood served as positive control; here we also deposited blood on denim pre-transfer (time 0) and post-transfer (24 hrs) were analyzed (Fig. 1; referred to herein as primary substrate).

**Figure 1.**
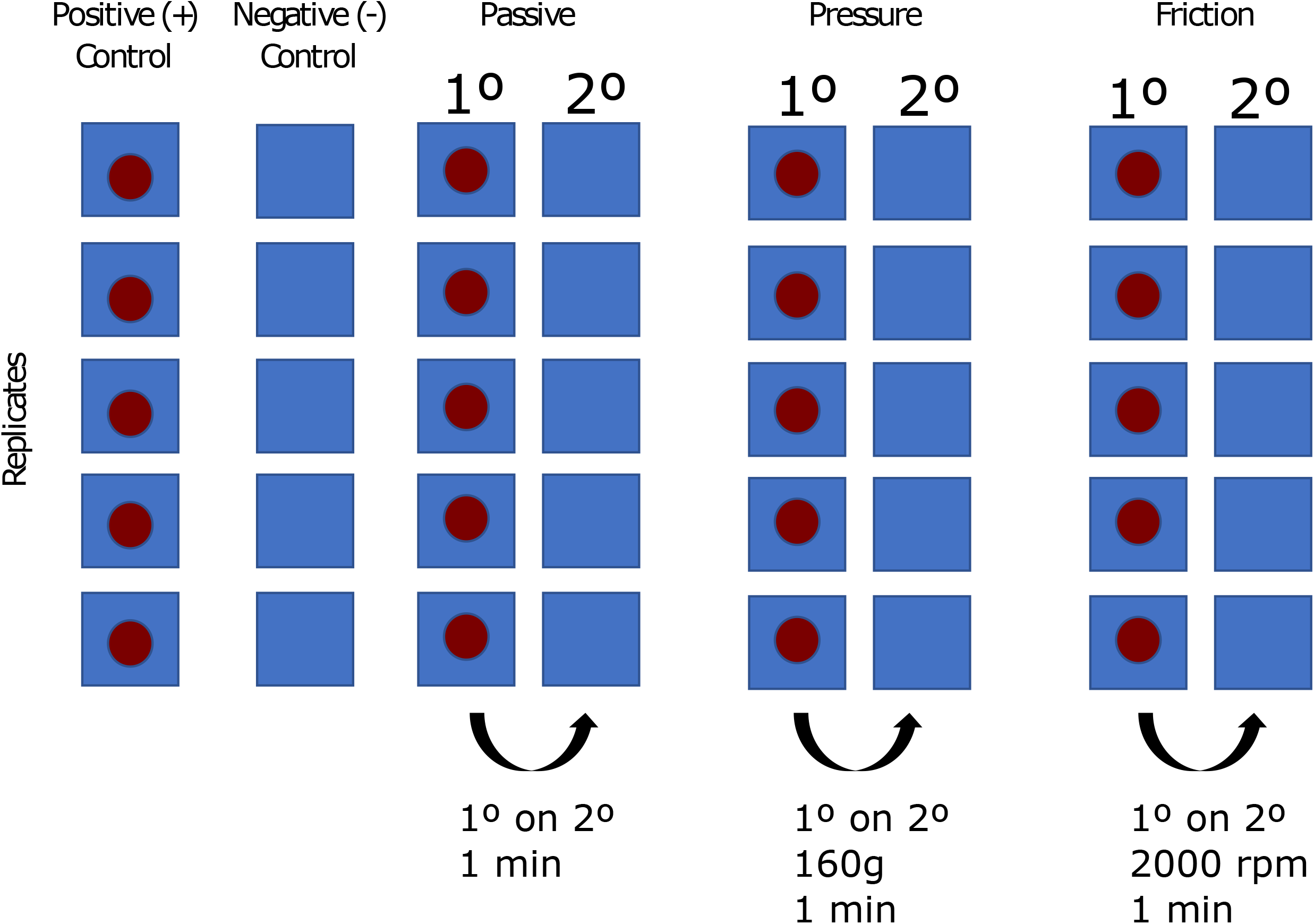
Experimental design for the controlled secondary DNA transfer event. Approximately 25 μL of fresh bovine blood was deposited on the primary substrate (A). After 10 minutes, the primary substrate (A) was placed on the secondary substrate (B) for 1 minute with three types of contact: passive, pressure, and friction. Positive and negative controls, where no secondary transfer occurs, were included. Substrates were 2×2 cm.

Approximately 24 hours after the transfer event occurred, we extracted DNA from the secondary substrates using the QIAamp DNA Investigator Kit (QIAGEN, Valencia, CA). The procedure from the QIAamp DNA Investigator Handbook for the *Isolation of Total DNA from Body Fluid Stains* was selected as it has a denim option to remove any inhibitors. Two modifications to increase DNA yield were applied: samples were centrifuged at 11000 rpm, rather than 8000 rpm, and the final incubation time for the samples at room temperature was five minutes, instead of one minute. The final elution volume was 50 μL. Transfer events were conducted separately for the qPCR and STR assays below.

### 2.3. DNA quantification and genotyping

The extracted DNA transfer samples were quantified using the Custom TaqMan Gene Expression Assay from Applied Biosystems and the QuantStudio 3 Real-Time PCR instrument. The primers were designed for specific amplification of the Cytochrome B gene of bovine mitochondria [27]. A qPCR standard calibration curve for was constructed using high-quality DNA extracted bovine tissue that was quantified on a Qubit fluorometric assay and diluted to set concentrations. The reaction mixture for each qPCR was made according to Supplemental TableS1. A no-template-control (NTC) sample containing no DNA was used as a negative qPCR control. Each extracted DNA sample was run in triplicate to account for pipetting error and calibrate the concentration estimates. The qPCR was set to 1 cycle of 95°C for 20 seconds, followed by 40 cycles of 95°C for 1 second, and 60°C for 20 seconds.

Bovine specific STRs were amplified using a primer mix that are a modified version of the StockMarks™ bovine primers (ThermoFisher Scientific; Table S2). PCR took place in a SimpliAmp thermal cycler following the conditions shown in Supplemental Table S3. After amplification, we added 1 μL of the PCR product to individual wells containing 9 μL of a solution HiDi formamide that included 500 Liz size standard, which was genotyped on the ABI 3730. STR PCRs and genotyping were replicated once for each sample. Negative controls (i.e. denim with no blood) were included.

### 2.4 Data analysis

The qPCR concentration was calculated as the average per sample from the triplicated assay: here, replicates with no amplification were assigned a concentration of 0 ng/μL. A Fisher Exact test was performed to test for the significance of type of contact on the success rate of DNA transfer: success was quantified as the simple presence of a qPCR amplification with an average quantity >0 ng/μL. A Kruskal-Wallis (KW) test of variance was used to assess the significance of the differences in the concentration of DNA transferred between the different types of contact. A Mann-Whitney-Wilcoxon (MWW) test was conducted to see which type of contact was significantly different than the others in a pair-wise fashion.

Scoring of the STR data was completed using Geneious Prime 2021.0.3 (https://www.geneious.com). Called peaks required a minimum RFU of 100 to remain conservative in the analysis [28]. Evaluation of genotypes was completed relative to the positive control – meaning the whole blood from the animal – referred to as reference; specifically, all transfer samples were compared to each locus of the reference. A transfer sample was given a score of 0 for no amplification or zero matches to the reference locus, 1 for matching one of the two reference alleles, and 2 for matching both reference alleles. For the latter, matching homozygous peaks were given a score of 2. This was done at each locus, and the sum of the scores across each locus was taken, providing an overall score for each sample relative to the control alleles (2 x no. of loci). We compared the genotype success between replicated contact types using a Pearson correlation and a paired t-test. As above, a KW test was first used to determine if there was a difference in genotype scores across the different contact types then a MWW test was completed to observe pairwise treatment differences.

All analyses and data visualizations were conducted in R Version 3.6.3 and, data and scripts are publicly available and deposited at: https://gitlab.com/WiDGeT_TrentU.

## 3. Results

DNA concentrations did not differ after 24hrs with addition of ACD-A (*t* = 0.86, *p* =0.38), suggesting similar transfer and degradation rates, so we merged experiments with and without ADC-A. All qPCR runs on the primary substrate amplified successfully (n = 31; 93/93 qPCR runs), but concentration decreased slightly upon transfer to the primary substrate both at time 0 and 24 hours. We observed no amplification in the qPCR and STR negative controls.

There were differences in the success rate of secondary DNA transfer between passive, pressure, and friction contact (*p* = 1.9e-06) with friction showing the highest transfer rate (Table 1; Figure 2). DNA was not detected in most passive contact secondary transfer events where >60% of qPCRs did not successfully amplify DNA. Friction contact had the highest concentration of DNA arising from secondary transfer (Figure 2). The primary substrate at time 0 and after 24 hrs did not statistically differ in concentration (*W* = 6, *p* = 0.07); but differed from all passive secondary transfer (*p* < 0.01). The MWW tests suggested no statistical difference between passive and pressure (*W* = 55.5, *p* = 0.57), but differences between friction and passive (*W* = 107, *p* < 0.01), and friction and pressure (*W* = 83.5, *p* = 0.01).

**Table 1:** Summary statistics for % qPCR amplification and % genotyping completeness for each type of contact and primary and secondary substrates. Technical replicates are shown in the *n* column. Mean qPCR concentration in ng /ul and ± standard deviation shown in parentheses. STR genotyping replicates 1 and 2 are separated by |.

| qPCR Assay |  | *n* | qPCR amplification |
| --- | --- | --- | --- |
| 1° | Denim | 6 | 100% (0.11±0.06) |
| | Denim - 24 hr | 6 | 100% (0.04±0.02) |
| 2° | Passive | 13 | 36% (0.03±0.01) |
| | Pressure | 10 | 70% (0.06±0.03) |
| | Friction | 10 | 93% (0.06±0.03) |
| STR Assay |  | *n* | Genotype completeness |
| 1° | Denim | 6 | 97% |
|  | Denim - 24 hr | 6 | 38% |
| 2° | Passive | 13 | 6% \| 30% |
|  | Pressure | 13 | 29% \| 32% |
|  | Friction | 13 | 40% \| 46% |

**Figure 2.**
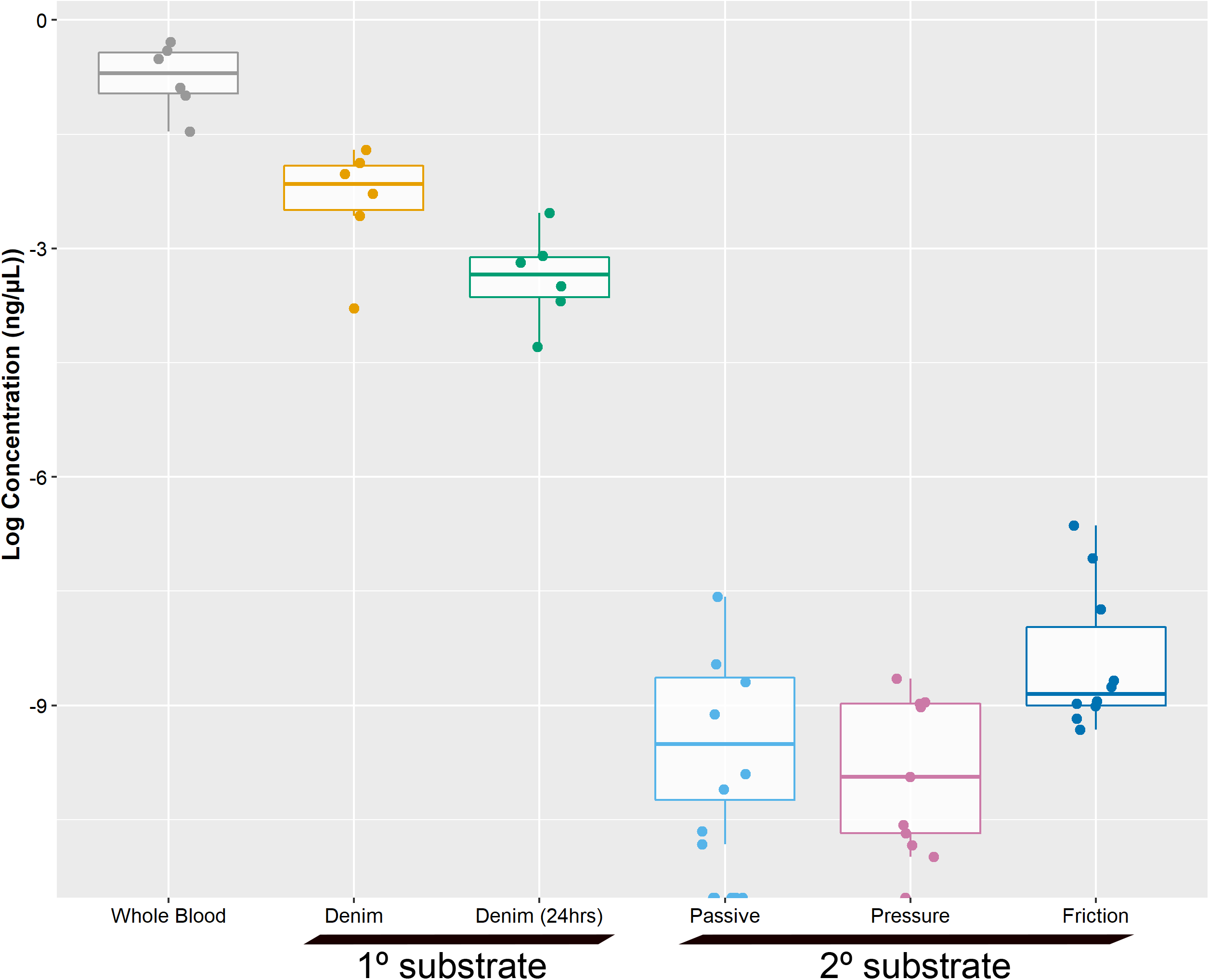
Concentrations (ng/μL) of DNA from the controls and the secondary substrate.

Genotypes were readily reproducible on the primary substrate but were diminished after 24 hrs (Table 1). The correlation of STR genotyping success between the two replicates by treatment was: passive (Pearson *r* = −0.28, *p* = 0.37), pressure (Pearson *r* = 0.45, *p* = 0.12), and friction (Pearson *r* = 0.67, *p* = 0.01). Trends were consistent between replicates (Figure 3) with pair-wise t-tests showing minimal differences (*p* > 0.05). The genotyping success showed high levels of variance among technical replicates (Figure 3). Transfer via friction provided the greatest resolution in terms of mean number of matching alleles; individual replicates were <50% complete (Table 1) but when we combined replicates genotypes were 59% complete for friction compared to 44% and 36% for pressure and passive respectively. The KW suggested a statistical difference in % genotype success for STR replicate 1 (*χ*^2^ = 13.02, df = 4, *p* = 0.01), but not replicate 2 (*χ*^2^ = 5.30, df = 4, *p* = 0.26). Differences in replicate 1 were driven by passive and friction comparisons (*W* = 26.2, *p* < 0.01) as the other comparisons had similar distributions (*p* > 0.05).

**Figure 3.**
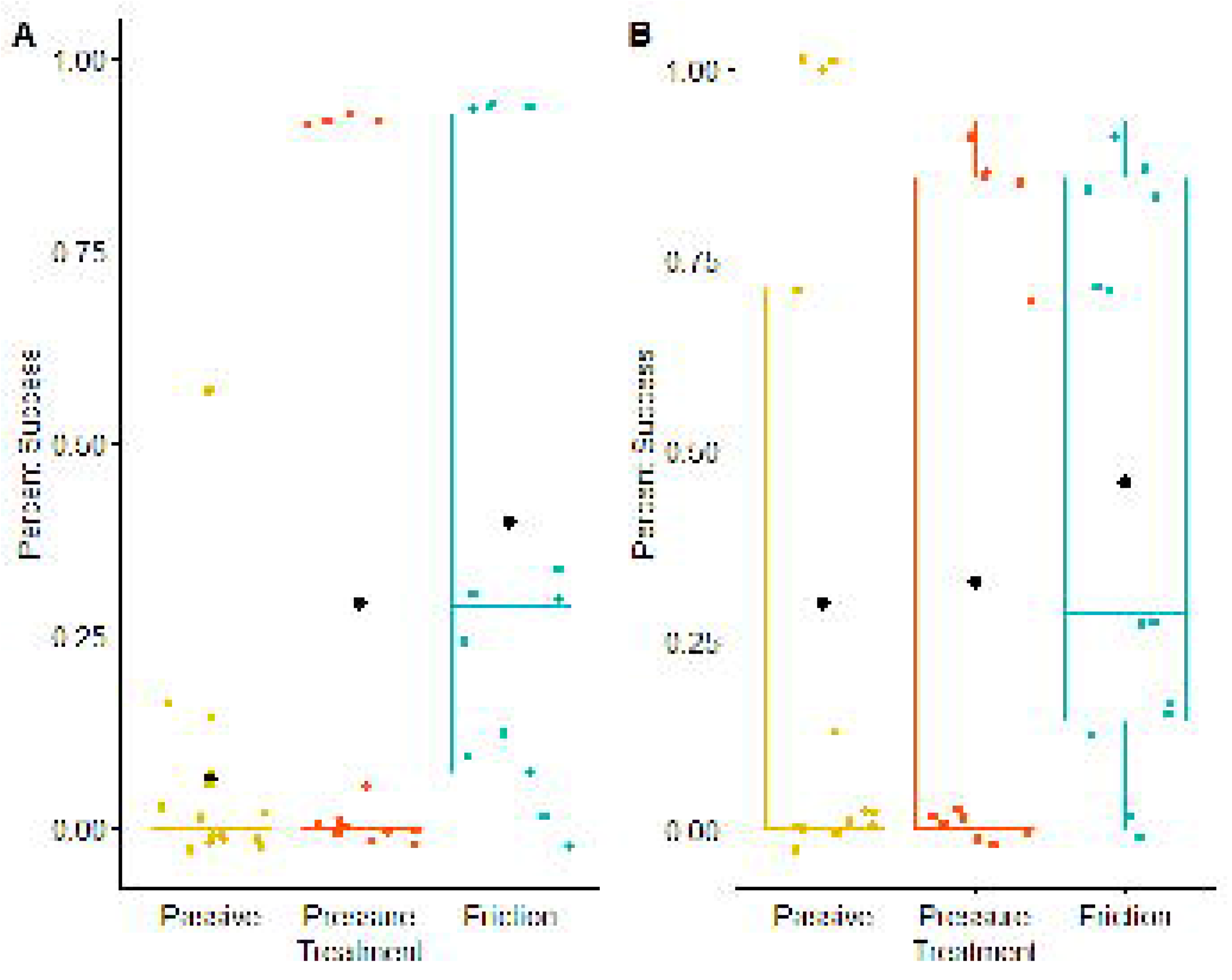
Percent successful genotype success by replicate 1 (A) and 2 (B). Black dots denote mean value, while horizontal bars represent the median.

## 4. Discussion

Secondary DNA transfer is more dependent on the nature of material and type of contact than the biological source [12]. Friction, or the act of two surfaces rubbing against each other, has shown to be conducive to DNA transfer [12]. While differences in the amount DNA transfer were documented in our experiment, this resulted in stochastic amplification and minor trends as it pertained to the STR profiles and contact type, with less than 50% genotyping success on average [29]. The inability to produce full STR profiles is best explained by the low template (LT) and degraded DNA on the secondary substrate, regardless of contact type. Thus, even though LT DNA is readily transferred, full profiles were difficult to generate, limiting their forensic utility or suggesting that adjusted laboratory protocols might be required. The first or primary transfer does also appear to be a factor, as DNA concentration was reduced, and notably genotype completeness decreased after 24 hrs; thus, isolating DNA from a transfer event immediately might improve the genotype, however unlikely this is to be in actual casework.

In LT scenarios, mtDNA generally amplifies better because of the high copy number [30]; in contrast, nuclear DNA often shows stochastic amplification patterns [31] and allelic drop-out [32]. While more DNA was transferred between denim under friction scenarios (Figure 2) differences in STR success were less pronounced (Figure 3). Our low to moderate STR success rate was quite similar to other STR methods for LT and secondary transfer DNA experiments [29,33]. While partial profiles might still be useful [34], it is generally accepted to repeat PCRs to iteratively build profiles [35]. Benschop et al. [29] suggested four PCR replicates was ideal for LT DNA, with both increased capillary electrophoresis injection and number of cycles improving genotyping success. Analytical approaches to optimize profile building from replicates [29] and modify likelihood ratio tests [36] can also be applied in such scenarios. While a validated protocol and a consensus has yet to emerge^1^, it was clear secondary DNA transfer yielded suboptimal profiles in our experiment that would limit their utility. Given the similarities between human and cow blood [26] and genomes [37], and the commercial STR assay used, we expect similar patterns to be present in human blood. We also reiterate that we observed little to no heme and indigo inhibition as there was only a slight

The growing interest in transfer and trace DNA (e.g [10,13]) has led to interest in experimental approaches [14]. Experiments have often focussed on substrate, contact type, and biological material [12,16], and combined with our study can identify patterns that inform opinions as it pertains to DNA transfer. It is true studies such as ours establish valuable baseline data [15], but arguably the most important pattern might be the ubiquity of secondary transfer. Akin to how forensic genealogists will regularly find a relative to a sample in a DNA database [38], one will often find [secondary] trace DNA if looking for it.

## Acknowledgements

The authors would also like to thank Otonabee Meat Packers for providing the blood samples. The Natural Resources DNA Profiling & Forensics Centre (NRDPFC) provided primers for this project. Thanks to Dr. Theresa Stotesbury and Prof. Rhonda Smith for providing comments. This work was supported by an NSERC Discovery Grant to Aaron B.A. Shafer (RGPIN-2017-03934) and a CFI-JELF (#36905) award to Aaron B.A. Shafer. Colin Elliot and Rebecca Ridings were supported by a NSERC-USRAs. Thanks to Reni Verasztó for help with genotyping.

## Footnotes

1 https://strbase.nist.gov/LTDNA.htm

