## Supplemental Tables 1-3 for "Secondary DNA transfer on denim using a human blood analogue"

**Table S1**: qPCR reagents and volume for one sample

| **Volume (μL)** | **Reagent** |
| --- | --- |
| 10 | TaqMan Multiplex Master Mix |
| 1 | TaqMan Gene Expression Assay |
| 7 | DNase/RNase-free water |
| 2 | DNA |

**Table S2**: Primers included in PCR.

| Locus | Dye | Colour | Expected Size Range (bp) |
| --- | --- | --- | --- |
| TGLA227 | FAM | Blue | 64 – 115 |
| ETH10 | FAM | Blue | 198 – 234 |
| SPS115 | FAM | Blue | 235 – 265 |
| TGLA126 | HEX | Green | 104 – 131 |
| INRA23 | HEX | Green | 193 – 235 |
| ETH3 | NED | Yellow | 90 – 135 |
| ETH225 | NED | Yellow | 136 – 165 |
| BM1824 | NED | Yellow | 170 – 218 |

**Table S3**: Thermal-cycling conditions for eleven-plex PCR

| Times and Temperatures | | | | | | |
| --- | --- | --- | --- | --- | --- | --- |
| Initial  Step | Each of 31 Cycles | | | Final  Extension | Final  Step | Hold  at |
|  | Melt | Anneal | Extend |  |  |  |
| 1 cycle  95°C  10 min | 94°C  45 sec | 61°C  45 sec | 72°C  60 sec | 1 cycle  72°C  60 min | 1 cycle  25°C  2 hours | 4°C  HOLD |
